# *Agrobacterium* mediated transformation and deciphering SNPs in *TaLr67* gene homeologs for gene editing in wheat

**DOI:** 10.1101/2022.03.23.485492

**Authors:** Karthikeyan Thiyagarajan, Luis Miguel Noguera, Mario Pacheco, Velu Govindan, Prashant Vikram

## Abstract

Diseases adversely affect grain yield of crop plants. Leaf rust is a major disease of wheat. As race-specific resistance breaks down, introduction of newer sources of resistance often from older accessions is necessary. Linkage drag from the donor accessions adversely impacts grain yield. As a result, CIMMYT breeding effort has shifted to using durable resistance, also referred to as race non-specific host resistance, which allows slow rusting but maintains grain yield. One of the key genes for durable resistance is *Lr67* and the resistant form of this gene is absent in CIMMYT elite lines. Hence, we have initiated efforts to convert the susceptible copy of *Lr67* into its resistant form directly in these elite lines using gene editing. This would eliminate backcrossing and thus save time as well as eliminate linkage drag that would accompany the resistant copy of the gene if it were to be introgressed from an older accession. As first steps, we have isolated and sequenced the genomic copies from each of the A, B, and D genome of the *Lr67* gene from three elite lines and an experimental line. Identification of more than 50 single nucleotide polymorphisms (SNPs) in the open reading frames (ORF) among these lines would be useful in designing the guide RNA molecules with precision. Further, we have streamlined genetic transformation of these elite lines, a prerequisite step for gene editing.

## Introduction

Wheat provides approximately 20% of the calories to the world population on average (Hawkesford et al. 2013). Potential grain yield is the ability of the locally adapted crop varieties to produce grain under nonlimiting environmental conditions, for example, irrigation and fertilizers, and absence of disease and insect pressure (Wulff and Dhugga 2018). Whereas the potential grain yield of wheat has been reported to be ~13 t/ha, the actual yield is 3.5 t/ha. The harvested yield is so much lower because of limitation in growth that results from limited water or fertilizer availability plus disease and insect pressure. Competition from weeds and other unpredictable factors, for example, wind-induced lodging, further suppress the yielding potential of crop plants (Oerke 2006).

Farmers have benefited substantially from the availability of the genetically-modifies (GM) crops (Klumper and Qaim 2014). Further, bumper crop harvests have helped stabilize grain prices. The GM traits have been available and proven to be safe in maize and soybean for more than 20 years, but none has been approved for commercial planting in wheat (Klumper and Qaim 2014; Wulff and Dhugga 2018). GM crops have encountered resistance from the consumer groups, gene editing offers an alternative avenue to expedite varietal development (Voytas and Gao 2014). Development of new varieties by combining the conventional breeding and genetic engineering is an effective strategy for improving the agronomic and physiological traits in crops. Gene editing, which was first accomplished by oligo-mediated homologous recombination but with a very low rate of success, has evolved over time as attempts have focused on increasing the specificity and frequency of edited events (Carroll 2017). Nucleotide editing has been achieved with varying degrees of success using zinc fingers, transcription activator-like effector nucleases (TALENs), and mega nucleases (Carroll 2017). Recent introduction of a new technology, clustered regularly interspersed short palindromic repeats (CRISPR) and CRISPR-associated protein 9 (CRISPR-Cas9), has revolutionized the field of gene editing because it is easy to use and returns edited events in high frequency (Carroll 2017).

The conventional method to introduce new traits into cultivated varieties has been through crossing an elite variety to a donor line that contains the desirable trait, followed by repeated backcrossing to the elite line to reduce the genetic contribution of the donor line other than the gene of interest. Once disease resistance breaks down, however, newer sources of resistance must be introduced (Singh et al. 2016). Additional sources of resistance are generally found in wild relatives of the crop plants (Wulff and Dhugga 2018). Linkage drag is a term used to describe the adverse effect of the undesirable genes of the donor parent on the performance of the newly improved variety. It is nearly impossible to eliminate all the undesirable genes of the donor parent through recurrent backcrossing to the elite variety. Linkage drag can manifest in different ways, like increased root or stem lodging, which reduce grain yield. Gene editing offers an alternative to backcrossing if disease-susceptible elite lines can be directly transformed. Editing of the susceptible copy of a gene to its resistant version directly in elite lines not only eliminates linkage drag, but it also provides an elite disease-resistant variety that could be used as a donor of disease resistance to susceptible lines. Plant transformation, thus, is the first step in successful gene editing. We have overcome this hurdle by successfully transforming three elite wheat lines and an experimental line from CIMMYT. In this report, we briefly discuss our work in wheat transgenics based preliminary analysis with key findings, especially the sequence characterization of *Lr67* gene from CIMMYT elite lines for our efforts in gene editing.

## Materials and methods

Wheat genotypes were grown in the greenhouse with proper irrigation and photoperiod at 16/8 h with an ambient temperature at 26°C/16°C day/night temperature. Three-week-old seedlings were harvested, and genomic DNA extracted using the DNeasy Plant Mini Kit (Qiagen). Genomic PCR was performed using Q5 High Fidelity DNA polymerase (New England Biolabs) with 3’ end mutation-based genome specific oligonucleotide primers (Table 1). Oligonucleotide primers were synthesized by T4 Oligo (Irapuato, Mexico). PCR products were gel purified using the QIAquick gel extraction kit (Qiagen). PCR products were sequenced directly by Langebio-CINVESTAV Genomic Services (Irapuato, Mexico). Nucleic and amino acid sequences were aligned using Vector NTI.

**Table 1.**
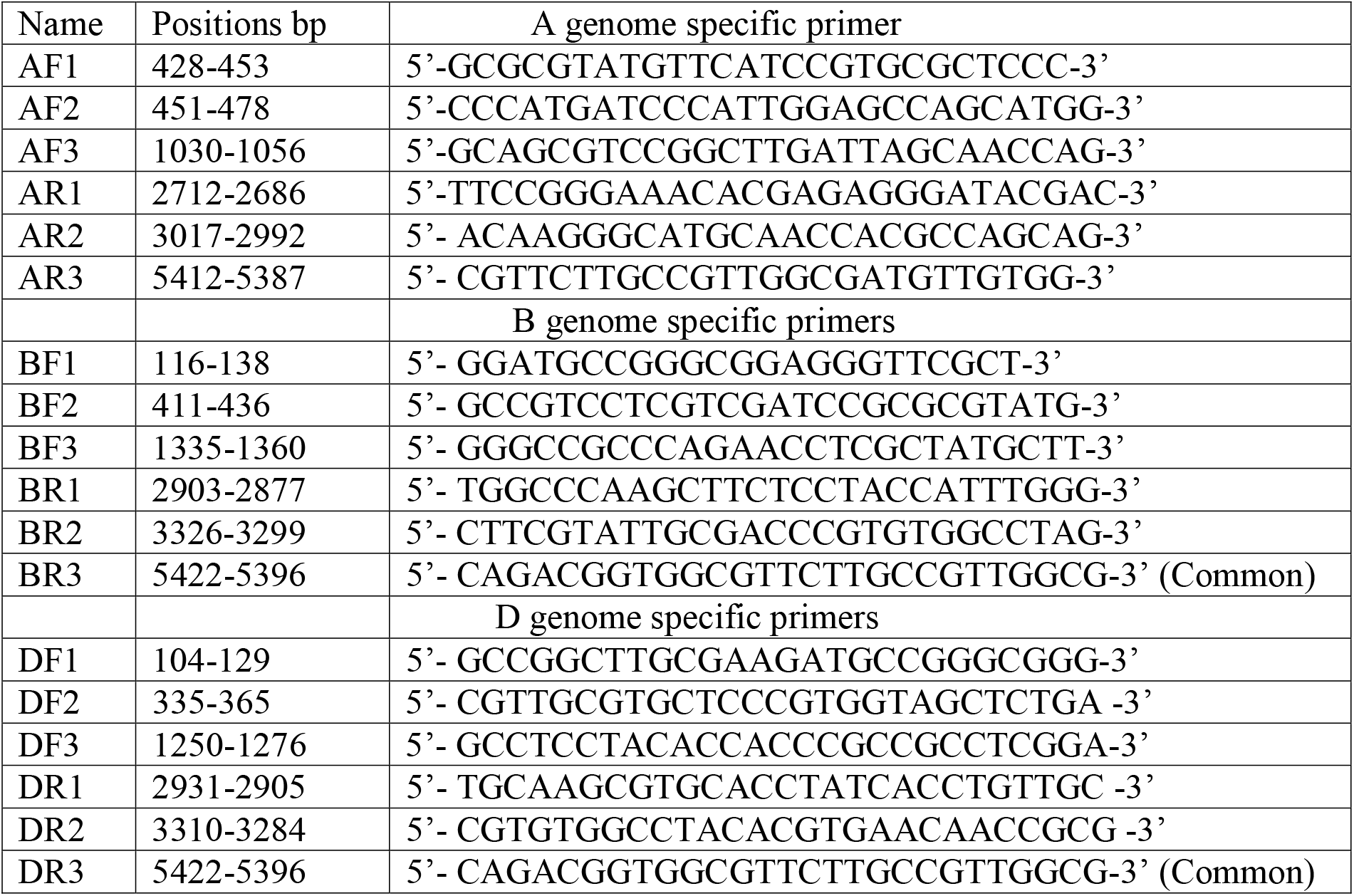
Amplification and sequencing of *Lr67* gene from A, B and D genomes, using 3’ end mutation-based genome specific primers.

Donor materials for explant preparation: At 15 DAF, spikes were collected, and immature seeds were extracted for each variety. The isolated seeds were sequentially surface sterilized in 70% (v/v) ethanol for 1 min, and 10% (v/v) Clorolex solution containing 100 μl of the surfactant Tween 80 for 10 minutes, successively three times rinsed with autoclaved distilled water to remove surface adhered sterilizing reagents. The embryos were excised under a light microscope.

Transformation was performed using *Agrobacterium tumefaciens* strain AGL1 (Lazo et al. 1991) using the protocol of Ishida et al. (Ishida et al. 2015) with some modifications as described. The AGL1 strain harbored a combinations of pAL154 and pAL156 vectors system derived from pSoup/pGreen constructs (Hellens et al. 2000), which were kindly provided by BBSRC, UK. The vector pAL154 possess 15kb Komari fragment, which transacts to induce the proliferation of pAL156 vector. The pGreen based plasmid PAL156 carries T-DNA integrating *Bar* gene (phosphinothrocin-resistant gene), *gusA* gene (modified β-glucuronidase gene) and maize RPOL intron inserted at 385th nucleotide to prevent the gene expression in *Agrobacterium* strain (Bourdon et al., 2001; Ke et al., 2002). Both genes were driven by ubiquitin promoter and ubiquitin intron derived from *Zea mays* (Christensen and Quail 1996). *Bar* and *gusA* genes were adjacent to left and right borders respectively. Purity of the *A. tumefaciens* AGL1 strain was maintained by adding 100 mg/l of kanamycin and 200mg/l of carbenicillin.

*A. tumefaciens* stock culture (430 μl) was inoculated in 10 ml of MG/L medium containing 20 μl of 100 mg/ml carbenicillin and 20 μl of 50 mg/ml kanamycin and incubated overnight at 37°C in a shaker at 250 rpm. Bacteria were quantified by measuring turbidity (OD at 0.4 at A600) after 16 h and the culture tubes were centrifuged at 3500 rpm (2000 g) in a centrifuge for 10 min.

Embryos were extracted from the immature seeds under a light microscope and the radicle excised with the help of scalpel and forceps. *A. tumefaciens* pellet obtained from centrifugation was briefly mixed with 4.8 ml 1X liquid media containing appropriate amount of silwet, acetosyringone and phloroglucinol. Isolated embryos (about 30 embryos for each microfuge tube) were immersed in 600 μl of the 1X inoculation medium containing *A. tumefaciens.* The tubes were mildly vortexed three times for 15-20 sec at 10 min intervals and incubated in a water bath for 3 min at 46°C. The treated embryos were transferred to the inoculation/co-cultivation solid medium. Subsequently, the induced embryos were transferred to the callus regeneration medium containing the antibiotic timentin.

The callus was then transferred to the induction medium and placed in a light chamber at 26°C for 4 to 5 weeks. Subsequent selection was done on medium containing phosphinothricin and timentin for 4 to 5 weeks and hardening of the plants was done in soil-filled mini pots for 4 to 5 weeks. GUS assays were performed as described (Wu et al. 2008).

Finale Bayer’s PPT, stock concentration 13.5 % was taken. Required of Ammonium Glufosinate concentration for PPT leaf painting assay was 2%, thus Finale Bayers PPT solution was diluted to obtain 2%, additionally 0.1 % Tween-20 was added in order to make an adhesive PPT leaf painting solution. Working solution preparation formulae: (13.5%/100) x 2%=0.27 ml or 270 μl of PPT was added to make 100 ml of aqueous solution, in which 0.1% or 100 μl of Tween-20 was also included. Matured leaves of both putative transgenic and non-transgenic Fielder varieties were painted using PPT solution upon dipping the cotton swab and painted over the selected area of the leaf tissues. Approximately after 3 to 4 days, the painted leaves of both transgenic and non-transgenic genotypes were observed for the leaf necrosis.

For Mendelian segregation analysis, about 9 to 30 matured seeds (T0) of T1 plants were harvested from green house and the kernels were cut transversely into two pieces. The seed with embryo was subjected for germination for further analysis, while remaining half part of the kernel was subjected for 12–24-hour GUS expression assay. The observed GUS positive to negative ratio was used for Chi-Square test to verify the Mendelian segregation ratio.

The wheat elite line Reedling was also transformed with a pCAMBIA vector, pSG19, where the expression of the DS-RED protein fused to a nuclear targeting sequence, N7, on the C-terminus, was driven by the CaMV 35S promoter and its transcription terminated by a NOS-polyA signal (Gillmor et al., 2010).

Selection was carried out on bialaphos, the resistance gene the expression of which was also driven by the CaMV 35S promoter and terminated by the NOS-polyA signal.

PCR volume was fixed with 2μl of dNTP (20mM each), 0.1 μl of Phusion Polymerase (2U/ μl) (Thermo Fisher Scientific Co.), 0.2 μl of 2 pM Forward primer, 0.2μl of 2 pM Reverse primer R, 0.6 μl of DMSO (100%), 4 μl of HF5X buffer, 0.5 μl of MgCl2 (50mM), 15.4 μl of Sigma H2O. PCR was performed using Mastercycler^®^ pro (Eppendorf) using the following cycling program: Initial denaturation at 98°C for 40 seconds, 35 cycles consist of 12 seconds denaturation at 98°C, 45 seconds annealing at 66°C, 23 seconds extension at 72°C and final extension at 72°C for 7 minutes. Primers used for *gusA* gene, forward: AGTGTACGTATCACCGTTTGTGTGAAC (Tm: 67.5°C), reverse: ATCGCCGCTTTGGACATACCATCCGTA (76.5°C). Primers used for *bar* gene, forward: GGTCTGCACCATCGTCAACC (58.6°C), reverse: GTCATGCCAGTTCCCGTGCT (Tm 60.4°C). Samples were stored at 4°C. Subsequently, PCR amplified fragments along with O’ Generuler 1kb ladder (Promega) were resolved using 1.2% of 10X diluted GelRed™ (Biotium Inc.) pre-stained agarose gel at 100 V for 2 h. The expected band was visualized using UV illumination-based documentation unit. Transgenic lines were confirmed with either *bar* or *gusA* gene.

## Results and discussion

### *Agrobacterium-mediated* transformation and transgenic screening

Editing native genes in the host genome is a promising approach to improve specific traits in crop plants. However, gene editing requires transformation of the host plant through biolistic or *Agrobacterium-mediated* methods. *Agrobacterium-mediated* transformation is the preferred strategy to edit genes in T0 generation as it results in simple, low copy integration of the T-DNA as compared to the biolistic method (Zhang et al, 2019). We have optimized the existing *Agrobacterium* transformation procedures and report here successful generation of transgenic events. (Fig. 1).

**Figure 1.**
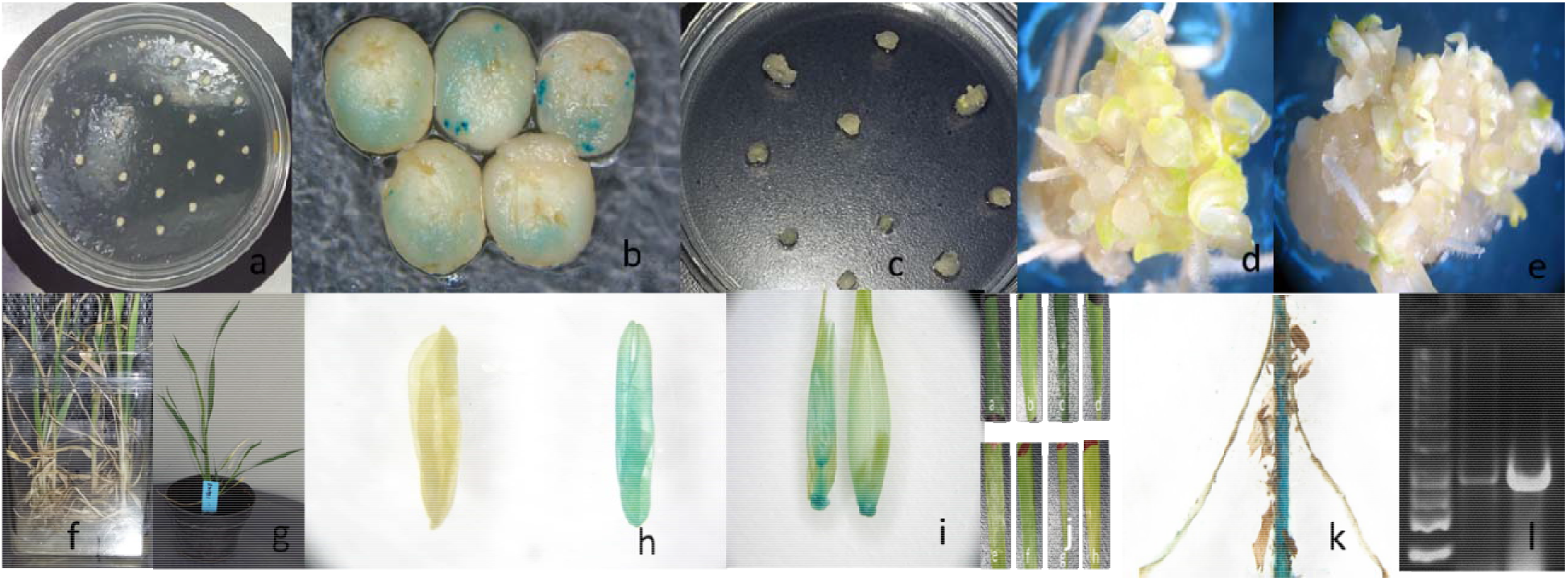
Schematic representation of wheat transformation. a) Infected embryos on growth induction media, b) transformed embryos expressing GUS, c) callus induction after 2 weeks, d) and e) regeneration of shoots after 3-4 weeks from pro-embryos of each callus in light chamber, f) PPT selection of young plantlets in light chamber, g) a PPT-selected young seedling, h) GUS assay showing difference between transgenic and non-transgenic anthers, i) GUS assay staining of young caryopsis, j) PPT leaf painting assay differentiating transgenic leaves without necrosis with non-transgenic leaves with necrosis, k) GUS staining of rootlet from transgenic plant, l) Transgenic confirmation with *Bar* gene fragment PCR amplification.

Callus bearing multiple shoots and roots from the regeneration medium was transferred to solid selection medium containing PPT. PPT inhibits glutamine synthetase in non-transgenic plantlets, which lack the *Pat/Bar* gene, causing senescence. Selected putative PPT-resistant seedlings were transferred to pots for hardening in a light chamber, where only the transgenic plants with apparently higher relative expression of the *PAT* or *BAR* gene survived. We observed that 2.5 mg/ml PPT stunted the plantlet growth along with root inhibition, and caused yellowing as previously described by Abdeen and Miki, 2009. Such senescence-prone plantlets with vestigial root formation along with retarded growth didn’t survive further in the light chamber, while healthy plantlets with resistance to PPT survived. PPT leaf painting assay showed that the transgenic genotypes exhibited a lack of necrosis or partial necrosis, apparently because of the presence of phosphinothricin acetyltransferase *(PAT)* or bialaphos resistance (*BAR*) genes (Fig. 2).

**Figure 2.**
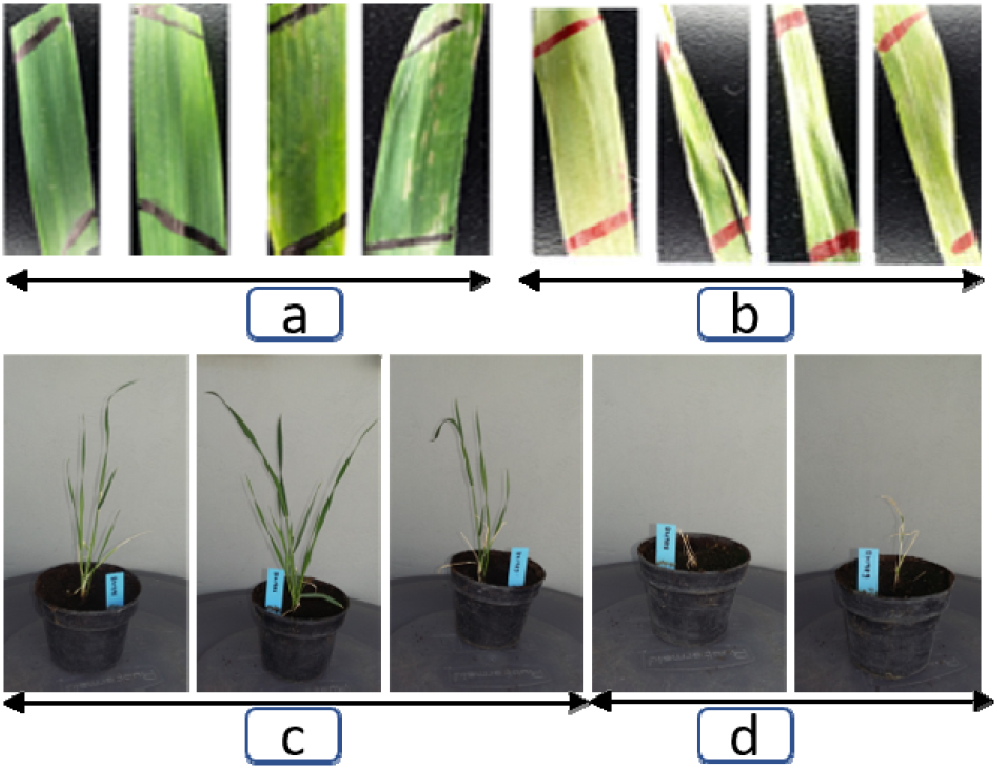
a) PPT painted leaves appeared as normal in putative transgenics, b) PPT painted leaves exhibited necrosis in non-transgenics, c) From left, the first three pots with healthy transgenic lines selected from PPT amalgamated media, d) Absence of apparent survival of last two genotypes with retarded growth from PPT selection media (experimental line Baj).

Successful transformation of Reedling, an elite CIMMYT variety, was carried out as described in M&M using a pCAMBIA vector containing the gene for the DS-RED protein fused at its C-terminus with a nuclear-targeting signal, N7, according to Gillmor et al. (2010). Targeting of the DS-RED fluorescent protein to nucleus is shown in Fig. 3, which demonstrates that we successfully transformed an elite CIMMYT variety.

**Figure 3.**
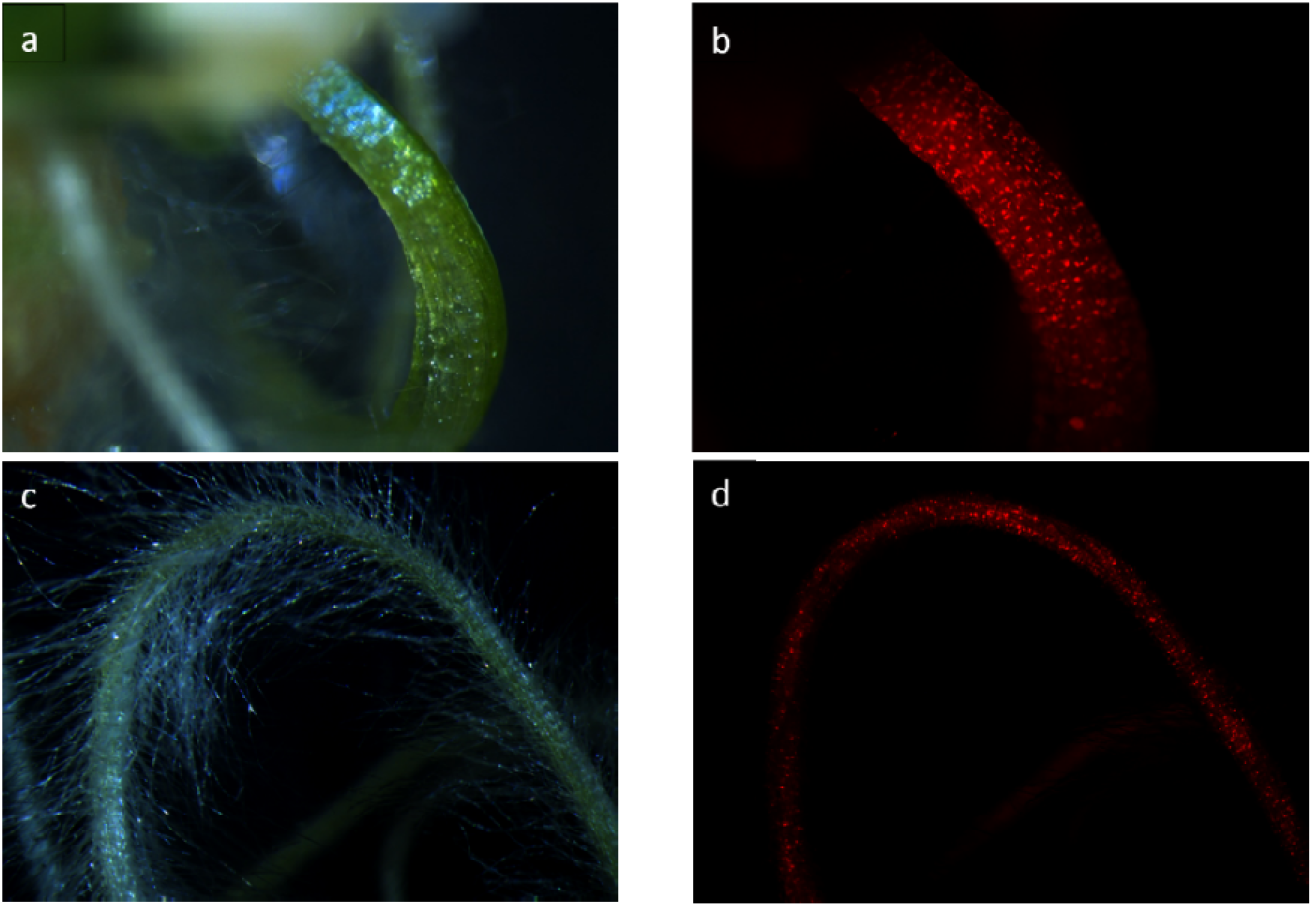
Transformation of the wheat variety Reedling with *Agrobacterium*. A vector expressing DS-RED driven by a 35S promoter was used to transform Reedling immature embryos. A regenerating seedling shoot (a) and root (c) visualized under visible light, and the same seedling shoot (b) and root (d) visualized under ultraviolet light using a DS-RED filter. DS-RED can be clearly seen in the nuclei, where it is targeted (right).

Other results are displayed in Fig. 4. Transformation efficiency ranged from 40-80 %. The presence of the transgene was confirmed by PCR of the *Bar* gene fragment size about 421 bp (Fig. 4). We have designed gRNA molecules for *Lr67* gene that are common to all four genotypes and are currently in the process of transforming the lines in collaboration with Corteva Agriscience (Lowe et al. 2018; Lowe et al. 2016). The regenerated events will be screened for gene edits as described previously (Savitashev et al. 2015).

**Figure 4.**
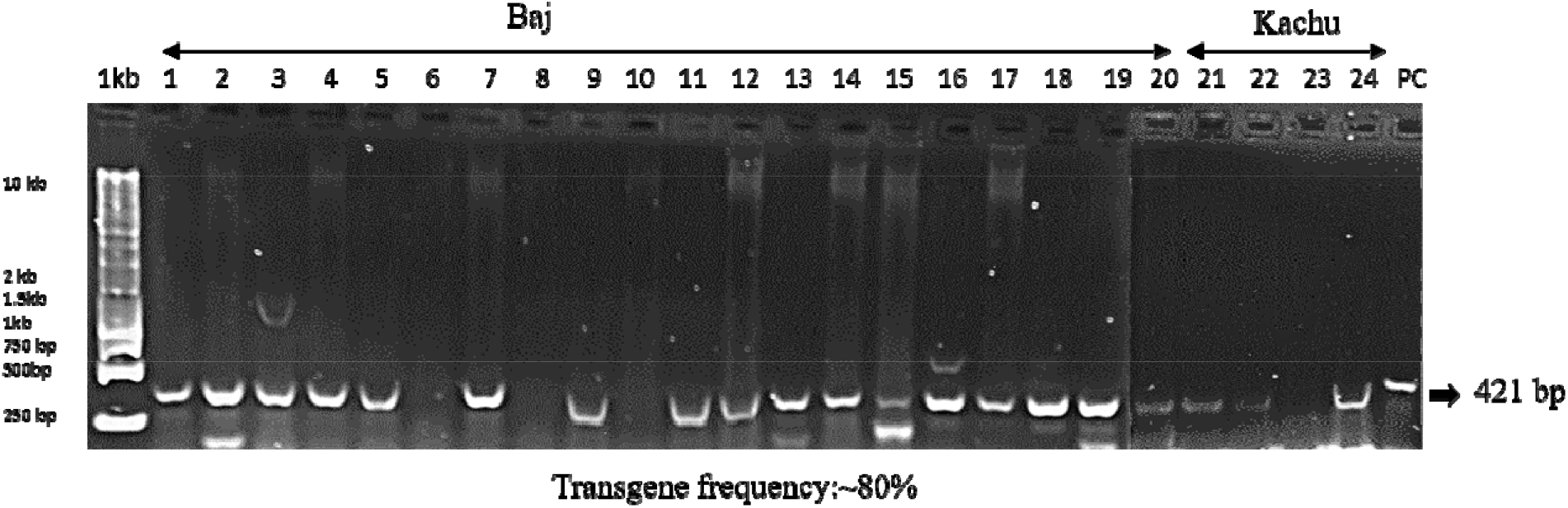
Amplification of *bar* gene in varieties Baj and Kachu. 1.1B, 2. 2B, 3.3B, 4.4B, 5.4B2. 6. 4B3, 7.5B, 8. 6B, 9.7B1, 10. 7B2, 118B1, 12.8B2, 13. 9B1, 14. 9B2, 15. 10B1, 16. 10B2, 17. 10B3, 18. 11B1, 19. 11B2, 20. 12B. Kachu genotypes: 21. KHA1, 22. KHB2, 23. KHB1, 24. KHC, 25. PC: Plasmid control.

### Mendelian Segregation analysis

The *GUS* assay based Mendelian inheritance test, resulted that expected positive to negative proportions were not completely comparable. However, the statistical significance states that there is a possibility of the comparison of expected and observed frequencies, mostly with 3:1 Mendelian-segregation ratio (Table 2) (Fig. 5). The reported range of non-mendelian segregation frequency between 10-50 % in plants (Yin et al., 2004), however, the transgenic progenies derived from *Agrobacterium* mediated transformation are usually complies with Mendelian inheritance (Budar et al., 1986) and in our analysis, phenotypic based segregation analysis showed the most T1 genotypes were in the pattern of Mendelian segregation.

**Table 2.** Phenotypic segregation analysis of seeds (T1) from T0 genotypes (variety Fielder) with Mendelian ratio tests. O (G+G-): Observed GUS Positive and Negative. E (G+G-): Expected GUS Positive and Negative. SR: Segregation Ratio. *P<0.05, **p<0.01, ***P<0.001.

| T<sub>1</sub> seeds of T<sub>0</sub>g | O (G+G-) | %T | E (G+G-) | SR | $\chi^2$ | P-Value |
| --- | --- | --- | --- | --- | --- | --- |
| T0_1_22seeds | 10:12 | 45 | 16.5:5.5 | 3:1 | 10.242 | 0.001372** |
| T0_1_22seeds | 10:12 |  | 11:11 | 1:1 | 0.182 | 0.67 |
| T0_2_9seeds | 1:8 | 11 | 6.75:2.25 | 3:1 | 19.59 | 9.58E-06*** |
| T0_2_9seeds | 1:8 |  | 4.5:4.5 | 1:1 | 5.444 | 0.02* |
| T0_3_30seeds | 1:29 | 3 | 22.5:7.5 | 3:1 | 82.178 | 1.2438E-19*** |
| T0_3_30seeds | 1:29 |  | 15:15 | 1:1 | 26.133 | 3.19E-7*** |
| T0_4_25seeds | 10:15 | 40 | 18.75:6.25 | 3:1 | 16.33 | 0.000053*** |
| T0_4_25seeds | 10:15 |  | 12.5:12.5 | 1:1 | 1 | 0.317 |
| T0_5_25seeds | 12:13 | 48 | 26.25:8.75 | 3:1 | 30.94 | 2.65738E-08*** |
| T0_5_5seeds | 12:13 |  | 12.5:12.5 | 1:1 | 0.04 | 0.841 |

**Figure 5.**
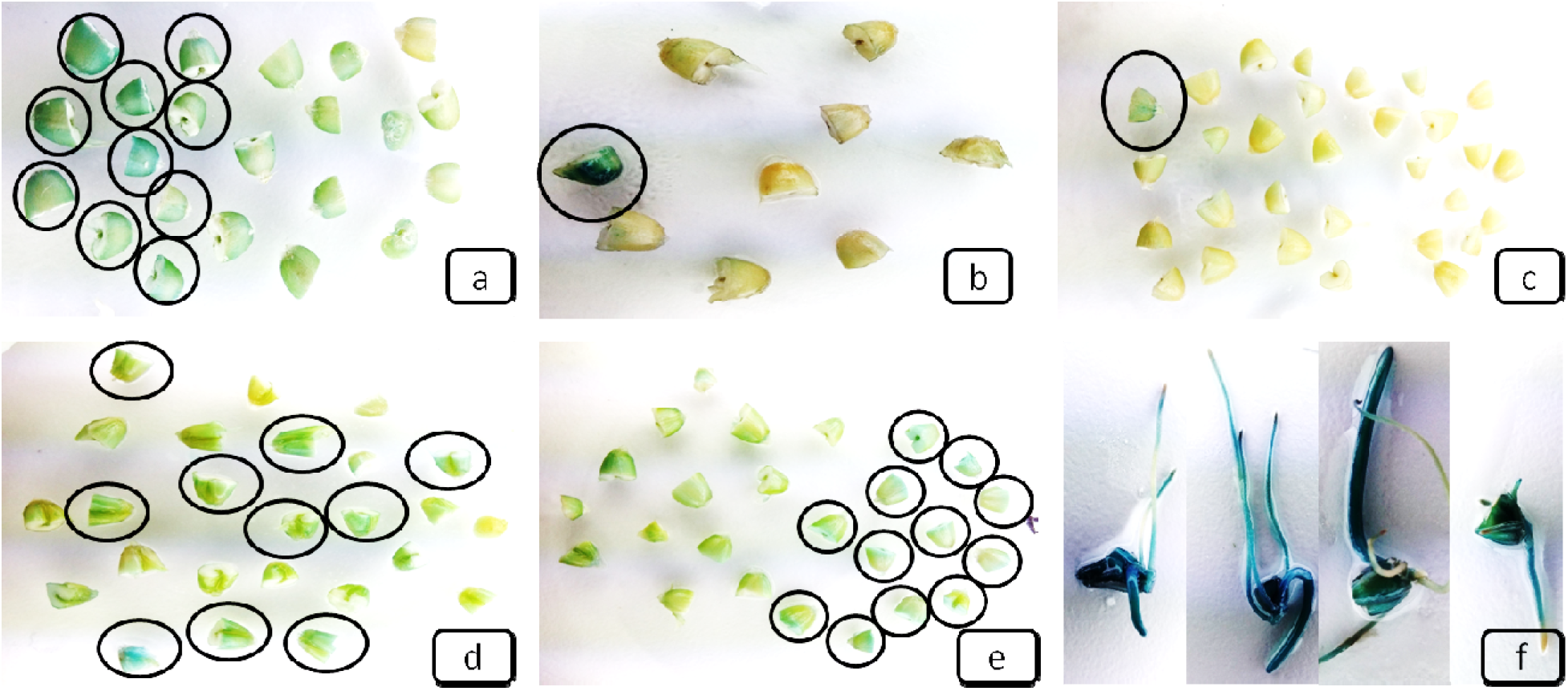
a-e) *GUS* staining-based segregation analysis of T_1_ seeds from T_0_ lines. *GUS* positive half seeds marked with circle. f) *GUS* positive seedlings.

### Sequence characterization of genomic copies of the *Lr67* gene for gene editing

*Lr67* hexose transporter gene is responsible for multiple disease resistance in wheat, and it has three exons and two introns (Moore et al 2015). The pairwise alignment of *Lr67* gene copies from A, B and D genome from Reedling with ENSEMBL gene models indicates the focused region with higher variation is in a copy from 4AS chromosome, while other two copies from 4BL and 4DL are relatively conserved than the copy from 4AS in the focused regions (Fig. 6).

**Figure 6.**
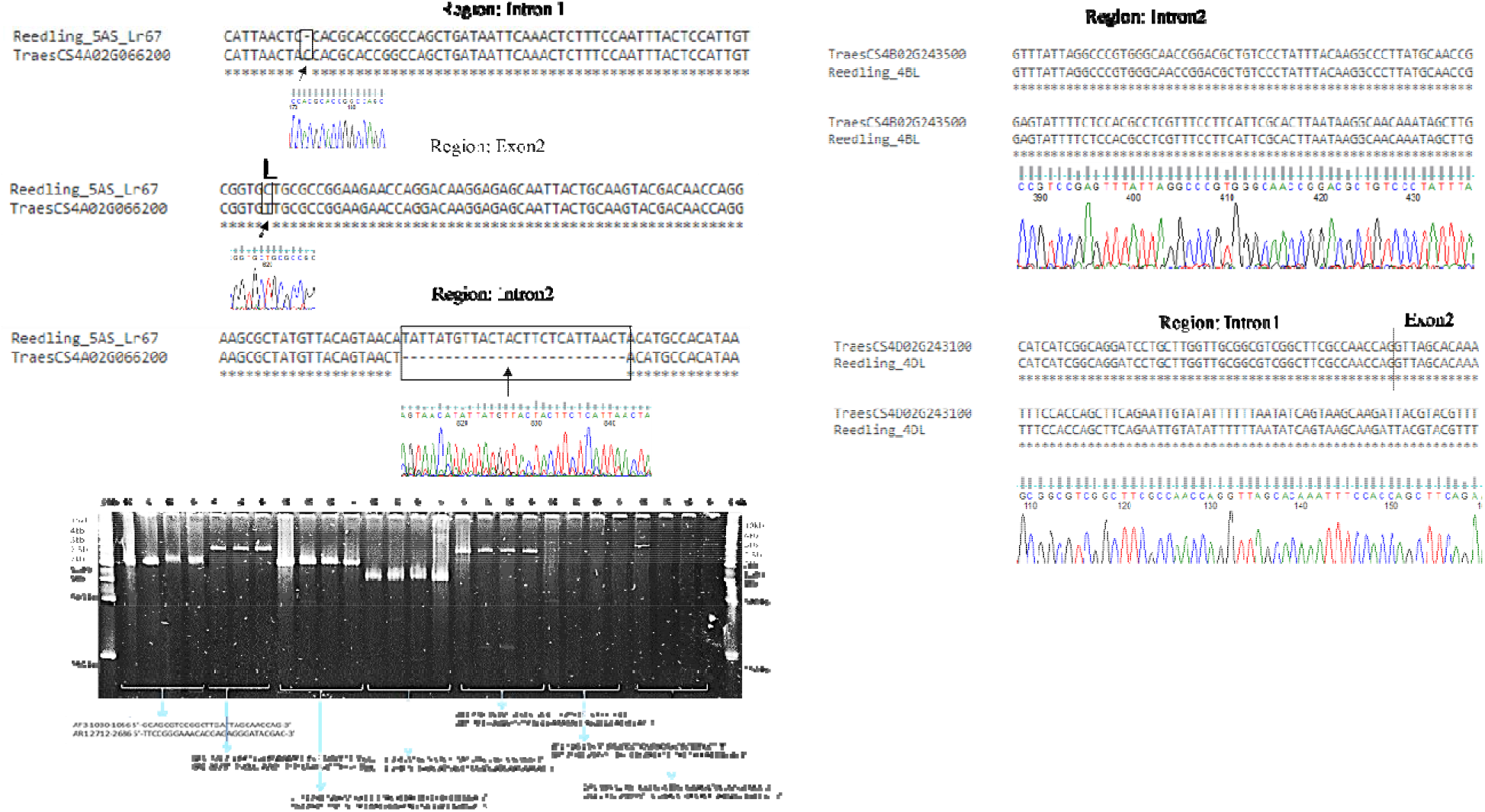
Amplification of *Lr67* gene fragments from corresponding genomes (4AS, 4BL and 4DL) of bread wheat varieties Reedling, Kachu, Baj and Fielder. Pairwise alignment of the *Lr67* gene fragments with existing ENSEMBL wheat gene model sequences and corresponding chromatogram regions from sequenced fragments.

Diseases reduce grain yield in wheat by approximately 13% on average (Singh et al. 2016). Leaf rust is the most widely adapted disease that causes varying levels of damage to the crop (Singh et al. 2016). We have specifically focused on *Lr67,* a gene that confers durable resistance against leaf rust but is only available in older accessions (Moore et al. 2015). In order to edit the susceptible copy of *Lr67* directly in elite wheat lines to its resistant form, we must first sequences each of the alleles corresponding to the A, B, and D genomes in the target genotypes. This step is necessary to avoid failure of the guide RNA to target the CRISPR-Cas complex to the gene of interest as even a single nucleotide mismatch could significantly weaken, if not eliminate, targeting. The genomic copies of *Lr67* were approximately 5 kb in length so we amplified allele-specific copies in parts. High quality Sanger sequence was provided by the DNA facility for approximately 700 nt for each run, requiring 7-10 sequence runs both for the forward and the reverse strands.

Two unique polymorphisms distinguish the resistant form of *Lr67* from its susceptible form (Moore et al. 2015). The first one is a G to C transversion at position 430 of the open reading frame (ORF), which causes a non-conservative change in the corresponding amino acid from Gly to Arg (Table 3, Fig. 7).

**Table 3.** Nonsynonymous polymorphisms in the open reading frame of wheat genotypes. Polymorphisms in the open reading frames of three elite CIMMYT lines, Baj, Kachu, and Reedling, an experimental line, Fielder, and the resistant (Res) and susceptible (Sus) open reading frames as reported in Moore et al. (2015) are presented.

| Codon (number) | Nucleotide (position in ORF) | Variant (amino acid) | Genotype |
| --- | --- | --- | --- |
| 144 | 430 | **CGT (Arg)** | **4D Res** |
|  | 430 | GGT (Gly) | 4D Sus, 4D |
|  | 430, 432 | GGC (Gly) | 4A, 4B |
| 151 | 453 | AAC (Asn) | 4D, 4B |
|  | 453 | AAG (Lys) | 4A |
| 238 | 713 | AAG (Lys) | 4D, 4B |
|  | 713 | AGG (Arg) | 4A |
| 358 | 1072 | TTC (Phe) | 4D, 4A, 4B Baj, 4B Fielder, 4B Kachu |
|  | 1072 | GTC (Val) | 4B Reedling |
| 375 | 1125 | AAG (Lys) | 4D, 4B |
|  | 1124 | AGG (Arg) | 4A |
| 376 | 1128 | TCG (Ser) | 4D, 4B Baj, 4B Fielder, 4B Kachu |
|  | 1128 | TCC (Ser) | 4A |
|  | 1127 | TTG (Leu) | 4B Reedling |
| 387 | **1161** | **TTG (Leu)** | **4D Res** |
|  | 1159 | GTG (Val) | All others |
| 408 | 1224 | CTC (Leu) | 4D Res, 4D Fielder, 4D Kachu, 4A, 4B |
|  | 1222 | GTC (Val) | 4D Sus, 4D Baj, 4D Reedling |
| 496 | 1488 | TTC (Phe) | 4D, 4B |
|  | 1487 | TAC (Tyr) | 4A |
| 502 | 1506 | CAC (His) | 4D, 4A |
|  | 1506 | CAA (Gln) | 4B |
| 504-505 | 1513-1515 | CAC insertion | 4A |

**Figure 7.**
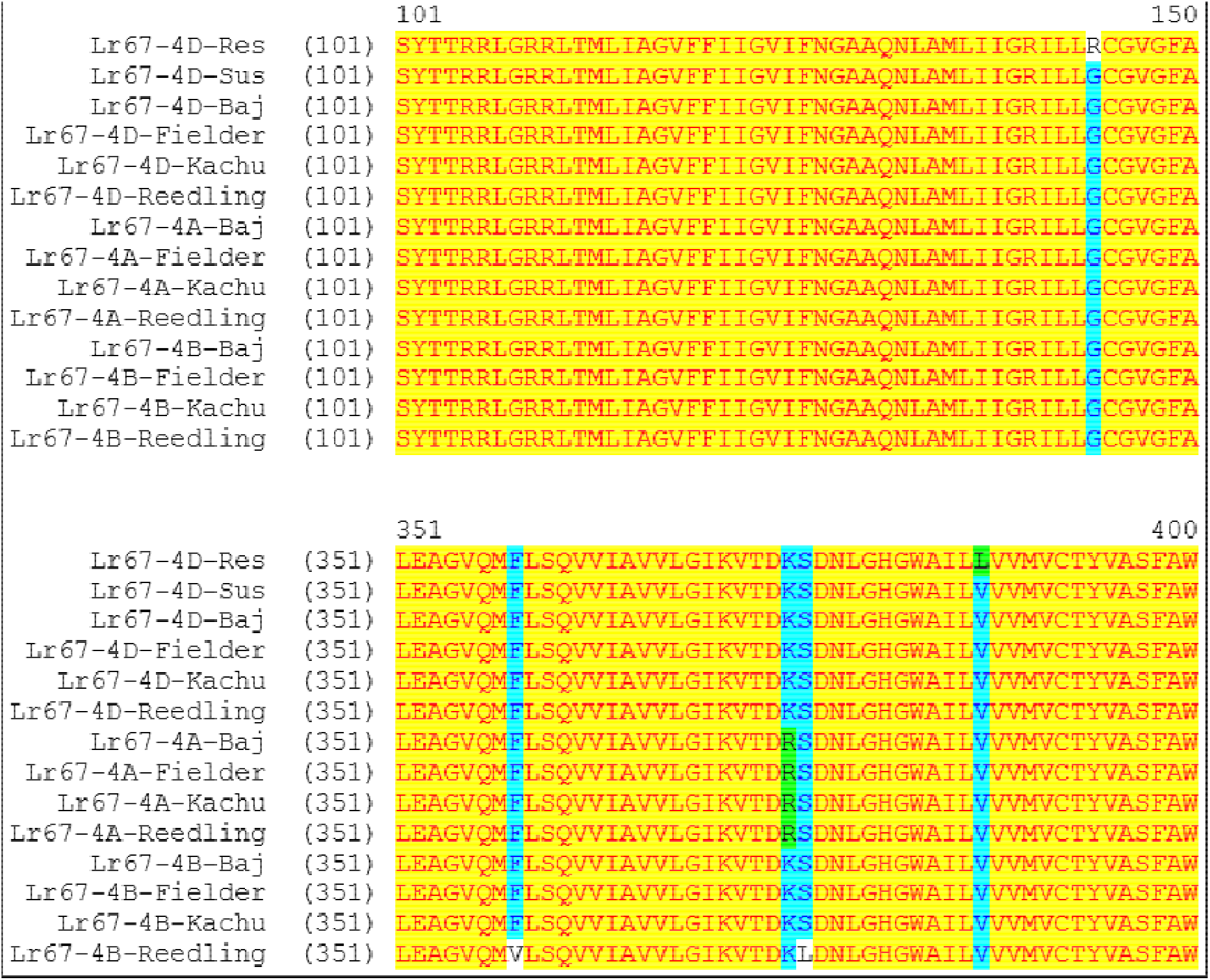
Sequence alignment of the Lr67 predicted protein for each of the genomes from four genotypes of wheat. The reference resistant (Res) and susceptible (Sus) alleles are included from Moore et al. (2015). Only the relevant portions of the protein containing the two polymorphisms (G144R and V387L) associated with rust resistance are shown.

The second is at nucleotide position 1161, G to T of the first nucleotide of the codon, which changes the corresponding amino acid from Val to Leu. This is a conservative change, however. Each of the polymorphisms corresponds to amino acids in a transmembrane domain (TMD). *LR67* protein is a hexose transporter and possesses 12 TMD (Moore et al. 2015). A non-conservative change like the one from Gly to Arg could cause a disruption in the TMD assembly and inactivate the protein (*LR67i*). Whereas Gly has only a hydrogen as a side chain and is neutral, Arg has a bulky, positively charged side chain and is an extremely basic amino acid with a pKa of 12.5, which would explain inactivation of the transporter protein. The second change, from Val to Leu, where both the amino acids are hydrophobic, is unlikely to disrupt the TMD hence is likely to have coevolved with the Gly to Arg mutation but without perhaps any role in the inactivation of the transporter protein. These observations were actually proved to be true using hexose uptake assays in yeast cells where the reversion of Arg to Gly in the encoded protein restored hexose uptake but Leu to Val did not when Arg (instead of the wildtype Gly) was present at the other site (Moore et al, 2015).

It is also likely that the transporter protein encoded by the *Lr67* gene, although annotated as a hexose transporter, is perhaps involved in the transport of toxin molecules of the pathogenic fungi into the plant cells. When it is inactivated, the remaining copies of the transporter encoded by the A and B genomes are unable to compensate for its loss. Our view is that it might require simple disruption of the *Lr67* ORF, which could be accomplished by employing the non-homologous end joining (NHEJ) after causing a cut with CRISPR-Cas, also referred to as the side-directed nuclease scenario 1 (SDN1) (Podevin et al. 2013). Further, by designing the gRNA molecules in the conserved regions of the ORF, it might be possible to disrupt the A and B copies as well, which will provide us additional tools to determine whether the inactivated *LR67i-A* and *LR67i-B* proteins phenocopy the *LR67i-D* protein, and if stacking them with the *LR67i-D* protein further augments rust resistance.

As to our objective of sequencing each of the *Lr67* copies from each of the genotypes to avoid failure of the gRNA to bind the target sequence, we identified 56 single nucleotide polymorphisms (SNPs) just in the ORF that is ~1.5 kb in length (only partial data shown in Table 1 and Fig. 5). Eleven of these polymorphisms led to nonsynonymous changes in the corresponding amino acids, eight of which are either non-conservative or semi-conservative in nature (Table 1). As discussed previously, the *LR67* protein from the resistant line is distinguishable from the susceptible line as well all the CIMMYT lines by two specific changes, a Gly144Arg and a Val387Leu where Gly and Val apparently correspond to susceptibility (Table 1 and Fig. 5). A unique change is the addition of a complete codon in the A genome near the end of the ORF (Table 1). Whether it has any significance is not clear.

We have initiated editing of the *Lr67* gene in Reedling using the tools developed in our laboratory. Successful identification of the edited events where the *Lr67* gene is inactivated would allow to test whether it confers additional resistance against a broad spectrum of fungi, something durable resistance genes are known to do (Moore et al., 2015). However, our experiments need further validation, hence we presumably conclude that our effort towards gene editing for accelerated breeding in wheat is a promising approach for developing new varieties with desirable features including durable disease resistance and yield improvement with precision.

## Acknowledgement

Authors are grateful to Dr. Kanwarpal Singh Dhugga (Head and Principal Scientist, CIMMYT Biotechnology) for his guidance and for helping to prepare the manuscript based on the data generated from CIMMYT Biotechnology laboratory. Authors are grateful to Dr. Kevin Pixely (Deputy Director General for Research (Breeding and Genetics), a.i., and Director of the Genetic Resources Program) and Ms. Rodelita Panergalin (Program Manager, Genetic Resources Program, CIMMYT) for their encouragement and support for this work.

## Author contributions

**KT**: Amplified, sequenced and assembled the genomic copies of *Lr67* gene, PCR transgenic confirmation, assisted and co-guided the tissue culture, helping to prepare the manuscript. **LN**: Tissue culture and *Agrobacterium* mediated transformation of Baj, Kachu, Fielder and Reedling wheat varieties. **MP**: Transformation of wheat line Reedling with a pCAMBIA vector pSG19 with the expression of the DS-RED protein. **GV** and **PV** provided critical suggestions and helped to improve the manuscript.

## Declaration

The authors have declared no competing interests.

## References

1. Abdeen A. and Miki B. 2009 The pleiotropic effects of the *bar* gene and glufosinate on the *Arabidopsis* transcriptome, Plant Biotechnology Journal. 7, pp. 266–282.

2. Budar F., Thia-Toong L., Van Montagu M., Hernalsteens J.P. 1986 *Agrobacterium-mediated* gene transfer results mainly in transgenic plants transmitting T-DNA as a single Mendelian factor. Genetics 114: 303–313.

3. Carroll D. 2017 Genome Editing: Past, Present, and Future. The Yale journal of biology and medicine 90 (4):653–659.

4. Christensen A. H and Quail P. H. 1996 Ubiquitin promoter-based vectors for high-level expression of selectable and/or screenable marker genes in monocotyledonous plants. Transgenic Research 5 (3):213–218. doi:10.1007/bf01969712.

5. Dhugga K.S. 2007 Maize biomass yield and composition for biofuels. Crop Science 47 (6):2211–2227. doi:10.2135/cropsci2007.05.0299.

6. Gillmor C.S., Park M.Y., Smith M.R., Pepitone R., Kerstetter R.A., Poethig R.S. 2010 The MED12-MED13 module of Mediator regulates the timing of embryo patterning in Arabidopsis. Development. 137(1): 113–22. doi: 10.1242/dev.043174.

7. Hawkesford M.J., Araus J.L., Park R., Calderini D., Miralles D., Shen T.M., Zhang J.P., Parry M.A.J. 2013 Prospects of doubling global wheat yields. Food and Energy Security 2 (1):34–48. doi:10.1002/fes3.15.

8. Hellens R.P., Edwards E.A., Leyland N.R., Bean S., Mullineaux P.M. 2000 pGreen: a versatile and flexible binary Ti vector for Agrobacterium-mediated plant transformation. Plant Molecular Biology 42 (6):819–832. doi:10.1023/a:1006496308160.

9. Ishida Y, Tsunashima M, Hiei Y, Komari T.2015 Transformation using immature embryos. In: Wang K (ed) Agrobacterium protocols, vol 1223. Methods in molecular biology, vol 1. Springer Science+Business Media, New York, pp 189–198.

10. Klumper W., Qaim M. 2014 A Meta-Analysis of the Impacts of Genetically Modified Crops. Plos One 9 (11). doi:e11162910.1371/journal.pone.0111629.

11. Lazo G. R., Stein P. A., Ludwig R. A. 1991 A DNA Transformation-Competent Arabidopsis Genomic Library in Agrobacterium. Bio-Technology 9 (10):963–967. doi:10.1038/nbt1091-963.

12. Lowe K., La Rota M., Hoerster G., Hastings C., Wang N., Chamberlin M., Wu E., Jones T., Gordon-Kamm W. 2018 Rapid genotype “independent” Zea mays L. (maize) transformation via direct somatic embryogenesis. In Vitro Cellular & Developmental Biology-Plant 54 (3):240–252. doi:10.1007/s11627-018-9905-2.

13. Lowe K., Wu E., Wang N., Hoerster G., Hastings C., Cho M. J., Scelonge C., Lenderts B., Chamberlin M., Cushatt J., Wang L. J., Ryan L., Khan T., Chow-Yiu J., Hua W., Yu M., Banh J., Bao Z. M., Brink K., Igo E., Rudrappa B., Shamseer P.M., Bruce W., Newman L., Shen B., Zheng P. Z., Bidney D., Falco C., Register J., Zhao Z. Y., Xu D. P., Jones T., Gordon-Kamm W. 2016 Morphogenic Regulators Baby boom and Wuschel Improve Monocot Transformation. Plant Cell 28 (9):1998–2015. doi:10.1105/tpc.16.00124

14. Moore J. W., Herrera-Foessel S., Lan C., Schnippenkoetter W., Ayliffe M., Huerta-Espino J., Lillemo M., Viccars L., Milne R., Periyannan S., Kong X., Spielmeyer W., Talbot M., Bariana H., Patrick J.W., Dodds P., Singh R., Lagudah, E. 2015 A recently evolved hexose transporter variant confers resistance to multiple pathogens in wheat. Nat Genet 47 (12):1494–1498. doi:10.1038/ng.3439

15. Oerke E. C. 2006 Crop losses to pests. Journal of Agricultural Science 144:31–43. doi:10.1017/s0021859605005708

16. Podevin N., Davies H. V., Hartung F., Nogue F., Casacuberta J. M. 2013 Site-directed nucleases: a paradigm shift in predictable, knowledge-based plant breeding. Trends in Biotechnology 31 (6):375–383. doi:10.1016/j.tibtech.2013.03.004

17. Savitashev S., Young J., Schwartz C., Gao H., Falco S.C., Cigan A.M. 2015 Targeted mutagenesis, precise gene editing and site-specific gene insertion in maize using Cal9 and guide RNA. Plant Physiology 169:931–945

18. Singh R. P., Singh P. K., Rutkoski J, Hodson D. P., He X. Y., Jorgensen L. N., Hovmoller M. S., Huerta-Espino J. 2016 Disease Impact on Wheat Yield Potential and Prospects of Genetic Control. In: Leach JE, Lindow S (eds) Annual Review of Phytopathology, Vol 54, vol 54. Annual Review of Phytopathology. Annual Reviews, Palo Alto, pp 303–322. doi:10.1146/annurev-phyto-080615-095835

19. Voytas D. F., Gao C. 2014 Precision Genome Engineering and Agriculture: Opportunities and Regulatory Challenges. Plos Biology 12 (6): e1001877. https://doi.org/10.1371/journal.pbio.1001877

20. Wu H. X., Doherty A., Jones H. D. 2008 Efficient and rapid Agrobacterium-mediated genetic transformation of durum wheat (Triticum turgidum L-var. durum) using additional virulence genes. Transgenic Research 17 (3):425–436. doi:10.1007/s11248-007-9116-9

21. Wulff B. B. H., Dhugga K. S. 2018. Wheat-the cereal abandoned by GM Genetic modification of wheat for disease resistance could help stabilize food production. Science 361 (6401):451–452. doi:10.1126/science.aat5119

22. Yin Z., Plader W., Malepszy S. 2004. Transgene inheritance in plants. J Appl Genet. 2004;45(2):127–44. PMID: 15131345.

23. Zhang Z., Hua L., Gupta A., Tricoli D., Edwards K. J., Yang B. and Li W. 2019. Development of an Agrobacterium-delivered CRISPR/Cas9 system for wheat genome editing, Plant Biotechnology Journal. 17, pp. 1623–1635, doi:10.1111/pbi.13088

